# Implications of the 375W mutation for HIV-1 tropism and vaccine development

**DOI:** 10.1101/2022.08.29.505747

**Authors:** Odette Verdejo-Torres, Tania Vargas-Pavia, Syeda Fatima, Paul R. Clapham, Maria J Duenas-Decamp

## Abstract

HIV-1 vaccines need to induce broadly neutralizing antibodies (bnAb) against conserved epitopes in the envelope glycoprotein (Env) to protect against diverse HIV-1 clades. To achieve this, we need to understand how different amino acids affect the Env trimer structure to find a common strategy to readily produce Env vaccines of different subtypes. Previously, using a saturation mutagenesis strategy we identified single Env substitutions that open the CD4bs without modifying the trimer apex. One of these substitutions was a tryptophan residue introduced at position 375. Here, we introduced 375W into a large panel of 27 T/F, acute stage, chronic infection, and AIDS M-tropic, and non-M-tropic primary isolates from clades A, B, C, D and G, and circulating recombinant forms (CRFs) (CRF02_AG, and CRF01_AE), and a complex (cpx) (CRF13_cpx). To understand the effect of 375W mutation on Env trimer structure and tropism, we evaluated soluble (sCD4) and monoclonal antibody (mAb) neutralization of wt and mutant Env+ pseudovirions using bnAbs (b6, 17b, b12, VCR01, 3BNC117, PGT128, 10-1074, PGT145, PG9 and PG16), as well as macrophage infection. Broadly neutralizing Abs (bnAbs) such VCR01, and 3BNC117 neutralized almost all the primary isolates tested while the other bnAbs neutralized many but not all of our panel. In general, 375W did not impair or abrogate neutralization of potent bnAbs. However, b12 and VCR01 showed some tendencies to neutralize 375W macrophage-tropic (mac-tropic) and intermediate mac-tropic mutants more efficiently compared with non-mac-tropic mutants. We identify wt and 375W mutant Envs in our panel that infected macrophages more efficiently than non-mac-tropic variants but did not reach the levels of highly macrophage-tropic brain reference Envs. These partial mac-tropic Envs were classified as intermediate mac-tropic variants. Surprisingly, we observed a mac-tropic (clade G) and intermediate mac-tropic (clade C, and D) primary isolates wt Envs that were not derived from the central nervous system (CNS). The 375W substitution increased sensitivity to sCD4 in all Envs of our panel and increased macrophage infection in many Envs tested including a CRF01_AE X4 variant. However, variants already highly mac-tropic were compromised indicating the presence of other factors implicated in mac-tropism. Increased sCD4 sensitivity and enhanced macrophage infection provide strong evidence that 375W confers exposure of the CD4bs across Envs from different clades/CRF/cpx and disease stages. Enhanced exposure of the CD4bs by 375W had little or no effect on exposure and sensitivity of CD4bs epitopes targeted by potent bnAbs. In summary, we show that 375W consistently increases Env binding to CD4 for diverse Envs from different clades and disease stages, 375W exposure of CD4 receptor is a biologically functional substitution that alone confers mac-tropism on non-mac-tropic Envs and 3) 375W is an ideal substitution for inclusion into HIV vaccines constructed from different subtype Envs, with the aim to elicit neutralizing antibodies that target the CD4bs while maintaining exposure of other Env broad neutralization sites, and 4) we found mac-tropic and intermediate mac-tropic Envs from blood indicating that these Envs could evolve outside of CNS or be released from Brain.

**Significance:** Substitutions exposing the CD4 binding site (CD4bs) on HIV-1 trimers, but still occluding non-neutralizing, immunogenic epitopes are desirable to develop HIV-1 vaccines. If such substitutions induce similar structural changes in trimers across diverse clades, they could be exploited in development of multi-clade Envelope vaccines. We show the 375W substitution increases CD4 affinity for Envelopes of all clades, circulating recombinant forms and complex Envs tested, independent of disease stage. Clade B and C Envs with an exposed CD4bs were described for macrophage-tropic strains from central nervous system (CNS). Here, we show that intermediate (clade C, and D) and macrophage-tropic (clade G) Envelopes can be detected outside CNS. Vaccines targeting the CD4bs will be particularly effective against such strains and CNS disease.

## Introduction

HIV uses the CD4 receptor and a coreceptor to infect cells. The main coreceptors are CCR5 and CXCR4. Primary isolates that infect via CCR5 (R5 viruses) were classified as non-syncytium-inducing or macrophage tropic (mac-tropic) because their envelope glycoproteins (Envs) were unable to induce syncytia in T-cell lines or peripheral blood mononuclear cells (PBMCs) and because they infected primary macrophages. More recently, highly mac-tropic R5 viruses or Envs were detected in samples from brain and CSF tissue, but so far mainly in subtype B and C primary isolates (1-4). A mac-tropic R5 Env was however detected in a semen sample from a patient infected by a group M virus which did not correspond to any described subtype or circulating recombinant form (CRF) (3). Moreover, highly mac-tropic Envs are able to infect target cells that express low levels of CD4 receptors such as human monocytes, microglial cells and hematopoietic progenitor cells as well as macrophages (5). Subsequently, non-macrophage tropic (non-mac-tropic) Envs that use CCR5 coreceptor were identified mainly in immune tissue, semen, and blood, and also in CSF (1, 2, 4).In contrast, variants that use the CXCR4 coreceptor (X4) are classified as T-tropic or syncytium-inducing due to their capacity to induce syncytia in T-cell lines and primary T-cells. Frequently, such viruses can use both CCR5 and CXCR4 coreceptors to infect cells and are termed dual tropic.

All attempts to develop a potent HIV-1 vaccine have been unsuccessful. The immunogens investigated so far have not induced potent broadly neutralizing antibodies (bnAbs) that are effective against each of the different HIV-1 subtypes and CRFs. An ideal immunogen candidate should recognize epitope structures common or similar across the huge range of Env variability of different HIV-1 clades. To achieve this, it is important that we improve our knowledge of how different amino acids and sites within Env impact the overall Env trimer structure and expose conserved epitopes such as the CD4 binding site (CD4bs) in the same way for all subtypes and CRFs. In fact, the CD4bs is a major target for highly potent, cross-reacting bnAbs e.g. 3BCN177 and VRC01 and also for the development of vaccines that aim to elicit neutralizing antibodies. It is therefore important to define sites and structures within the CD4bs that will need to be preserved in vaccines for the induction of neutralizing antibodies (nAbs).

However, it is important that Env immunogens do not divert the immune response to produce “off target” non-neutralizing Abs (non-nAbs) against occluded but highly immunogenic epitopes including CD4-induced (CD4i), V3 loop, or the trimer base of SOSIP immunogens (6). The glycan shield is used by HIV-1 to mask neutralizing epitopes enabling escape from immune system pressure. Removing a glycan on an immunogen surface can expose an epitope that will induce Abs and/or create novel immunodominant areas (7). However, such glycan holes are rarely present on different primary Envs. In addition, amino acid changes in Env may modify the trimer structure, shifting the orientation of proximal glycans and altering exposure of non-neutralizing epitopes (8).

Previously, we used EMPIRIC, a saturation mutagenesis assay to identify Env substitutions that opened, or exposed the CD4bs without major modifications to the trimer apex (9). The 375W substitution/mutant was one that was selected from a HIV-1 mutant library following cultivation in healthy donor peripheral blood mononuclear cells (PBMCs). In fact, position 375 was able to support a range of different amino acids that conferred replication as efficiently as (L, M, C, N, and Q) or better (H, T, Y, F and W) than wt (S) (9).

A tryptophan at position 375 in gp120 of YU2 strain that fills the Phe 43 cavity (10) has been implicated in slight increases of sCD4 affinity, 2G12 glycan monoclonal antibody (mAb) neutralization sensitivity, resistance to b12 mAb (against CD4bs) and other non-neutralizing CD4bs Abs (b6 and F105) (10, 11) as well as inducing a conformational change closer to a CD4-bound state (10-12) and refolding the bridging sheet (13). This 375W mutation can also be found in most HIV-2 and simian immunodeficiency (SIV) isolates such as SIVmac isolates including SIVmac239, SIV from sooty mangabey monkeys (SIVsmm) and some chimpanzee SIVs such as CPZ.TZ.06.TAN5, but it is not found in HIV-1 primary isolates (14). SIV can infect cells that express low levels of CD4 in the surface, and a tryptophan at position 375 could help explain why it is less dependent on the CD4 receptor (15). In SIV, substitution of the tryptophan for a serine (typically found in HIV-1) reduces CD4 interaction and infection. Other amino acid changes in layer 1 of gp120 also decrease CD4 affinity but this effect is counteracted by 375W mutation (15). SIV viruses that artificially carry HIV-1 Envs are called Simian-Human immunodeficiency viruses (SHIVs). HIV-1 strains cannot infect via rhesus CD4 (rhCD4). However, the introduction of basic (H) or hydrophobic (Y, F and W) amino acids at position 375 into clade A, B, C and D SHIVs not only increases infection in rhCD4 cells but also enables replication with titers close to HIV-1 in human cells without altering the sensitivity to bnAb (16).

The effect of other mutations at position 375 has been studied in the context of different subtypes. 375N in clade B HXB2 confers resistant to sCD4 and mAbs (39.13g, 1.5e, G13 and 448) that overlap the CD4bs (17). A substitution of serine for threonine, at position 375 confers resistance to BMS-626529 attachment, an inhibitor that binds HIV-1 Env to block CD4 interaction in subtype B (18). In a minority of CRF02_AG strains, histidine and methionine substitutions at 375 were genetically selected to induce drug resistance to this drug. CRF01_AE usually has histidine in position 375. When histidine is changed by a serine, viruses are less infectious (19) and a reduction in sensitivity to antibody dependent cellular-mediated cytotoxicity (ADCC) was also observed (20). Moreover, a tryptophan in position 375 in subtype B (YU2) and CRF01_AE reduced the gp120-gp41 association.

It is difficult to identify mutations that induce the same phenotype and/or conformational changes in the trimer for all clades including CRF and complex (cpx) Envs. In this study, we tested the effect of 375W on exposure of CD4bs and a range of epitopes targeted by bnAbs. The CD4bs and such neutralizing epitopes are the main targets for HIV-1 vaccines aimed at eliciting bnAbs. We tested a wide panel of diverse HIV-1 Envs in functional and neutralization assays and included transmitted/Founder (T/F), acute stage and late state mac-tropic and non-mac-tropic Envs of different subtypes such as clade A, B, C, D, and G, CRFs (CRF02-AG, CRF01-AE), and cpx variant (CRF13_cpx). In all Envs tested, the 375W substitution increased the exposure of the CD4bs as measured by sCD4 sensitivity as well as by their capacity to infect macrophages, with the exception of mac-tropic Envs that already carried an exposed CD4bs. Then, exposure of CD4 receptor via 375W is not the only factor responsible of mac-tropism. In summary, we describe for the first time, a substitution that increased sCD4 sensitivity and macrophage tropism in R5 Envs of all clades, CRFs and a cpx tested as well as a primary X4 Env. Using mAbs against other Env sites including CD4i (b17), V3 loop/glycan (10-1047, and PGT128) and trimer association domain (TAD) V2q (PG9, PG16 and PGT145), we show that 375W did not confer significant alterations to broadly neutralizing CD4bs epitopes on Env. Thus, 375W confers strong exposure of the CD4bs with only minor effects on V3 loop/glycan mAbs and other broadly neutralizing epitopes targeting gp120. However, the 375W substitution did alter the exposure epitopes for some less potent V1V2 loop/glycan mAbs. Finally, and of note, we also identified primary, mac-tropic (clade G) and intermediate mac-tropic (clade C, and D) Envs and that were not derived from central nervous systems (CNS). The role of such Envs in HIV pathogenesis is discussed.

## Results

### 375W mutation increases sCD4 sensitivity

To evaluate whether the 375W substitution influences soluble CD4 (sCD4) inhibition and therefore binding, we introduced 375W mutations by direct mutagenesis to a panel of Envs from different subtypes at different infection stages (see Materials and Methods, Table 1). T/F (3T, 6T, 15T, and 19T, and Z1792M), acute stage (Z153M, Z185F, Z221M, and clones_251, _269, _278 and _258), chronic infection (0503M02138), and AIDS (92UG975.10) M-tropic (B33, B59, JR-FL, B100), and non-M-tropic (LN40, LN8, JR-CSF and LN58) primary isolates. 3T, 6T, 15T, 19T, B33, B59, JR-FL, B100, LN40, LN8, JR-CSF and LN58 Envs are subtype B and Z153M, Z221M, Z185F, and Z1792M Envs are clade C. We also tested other primary isolates from clade A (BG505), clade D (NKU3006, A03349M1, A08483M1, 57128), clade G (92UG975), circulating recombinant forms (CRFs) such as CRF02_AG (Clone_251 and Clone_278), CRF01_AE (0503M02138, and Clone_269), and a complex (cpx) variant (CRF13_cpx Clone_258). The BG505 Env, that represented a mother and child transmission, was obtained 6 weeks after birth and the infant was infected at delivery or via breastfeeding (21) and used in vaccine development.

**Table 1:**
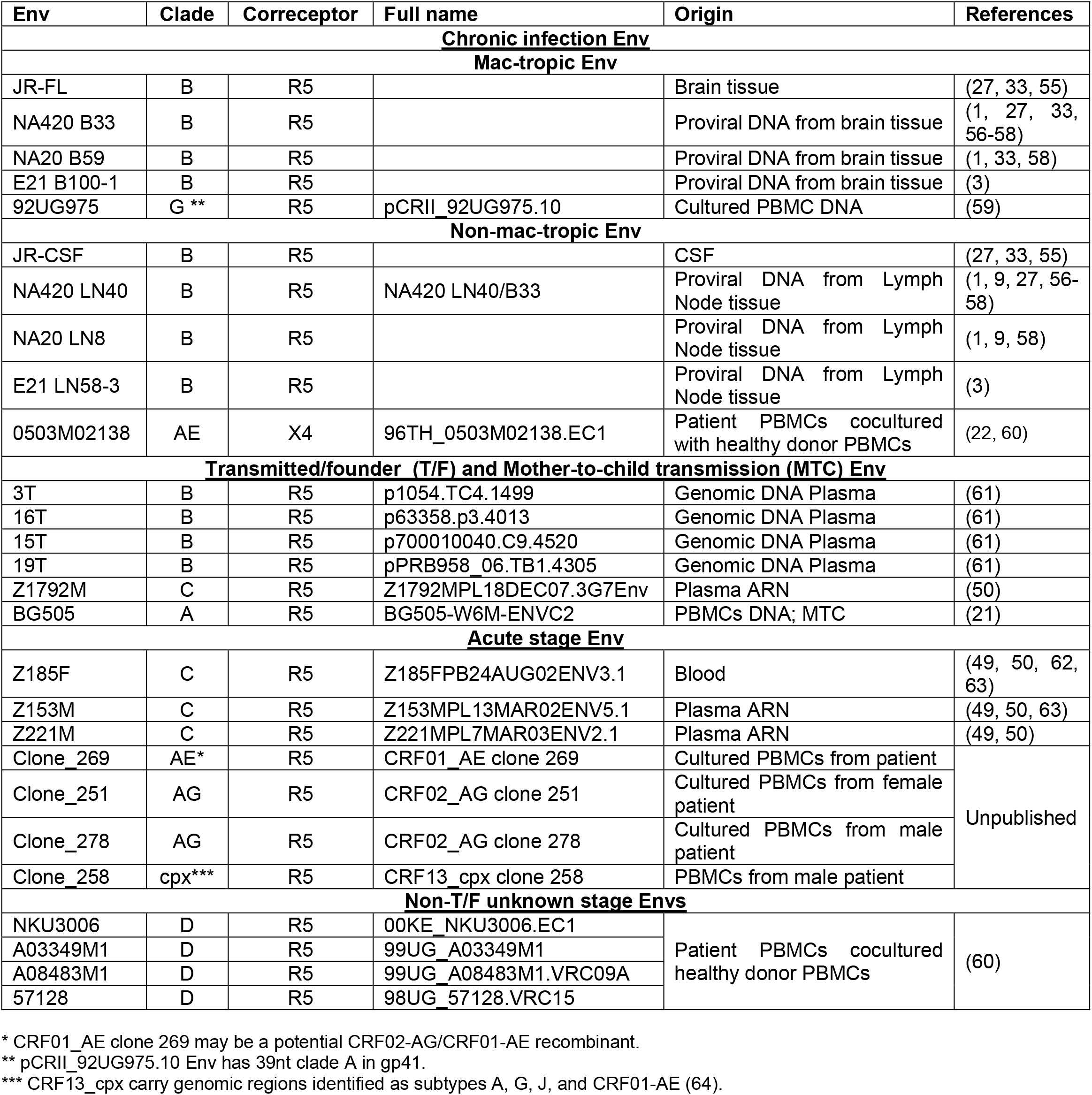
HIV-1 *env* clones used to prepare Env+ pseudovirions.

We first tested neutralization of all Envs starting at a maximum 50ug/ml of sCD4 and subsequent 2-fold dilutions. However, we could not calculate the IC50s of non-M-tropic LN8, LN40, LN58, JR-CSF, 6T, 19T, Z153M, Z185F, Z221M, Z1792M, BG505, clone_269, 0503M02138, NKU3006, A03349M1 and clone_278 Envs since there was less than 50% inhibition at the maximum amount of sCD4. We know that these envelopes can confer infection of cells, so they presumably bind to the CD4 receptor even though they must have low CD4 affinities. We decided to increase the sCD4 concentration up to 400ug/ml to accurately calculate IC50s of these Envs. In figure 1, we represent the IC50 along with 95% confidence intervals. We found, independently of clade or disease stage, that IC50 values are lower for pseudovirions carrying Envs with the 375W substitution showing that this mutation increases sensitivity to sCD4 neutralization. These data confirm our previous results that the presence of a tryptophan in position 375 opens the CD4bs facilitating the interaction with the CD4 receptor (9). Xiang et al. also reported an increase in sCD4 sensitivity for YU2, a subtype B isolate (10). In our experiment, there is at least a 1 log difference between most WT and mutant IC50s. B_B33, B_LN40, B_B59, B_LN58, B_JR-CSF, AE_0503M02138, and G_ 92UG975 mutants were exceptions and showed less than a log difference between wt and 375W mutant sensitivity to sCD4. Overall, we found significant changes in the IC50 values (*p* <0.001) when we compared all WT with all 375W mutant Envs (Figure 2). In summary, a tryptophan at position 375W increases sCD4 binding in all clades, CRFs and cpx tested, independently of disease state and compartment origin.

**Figure 1:**
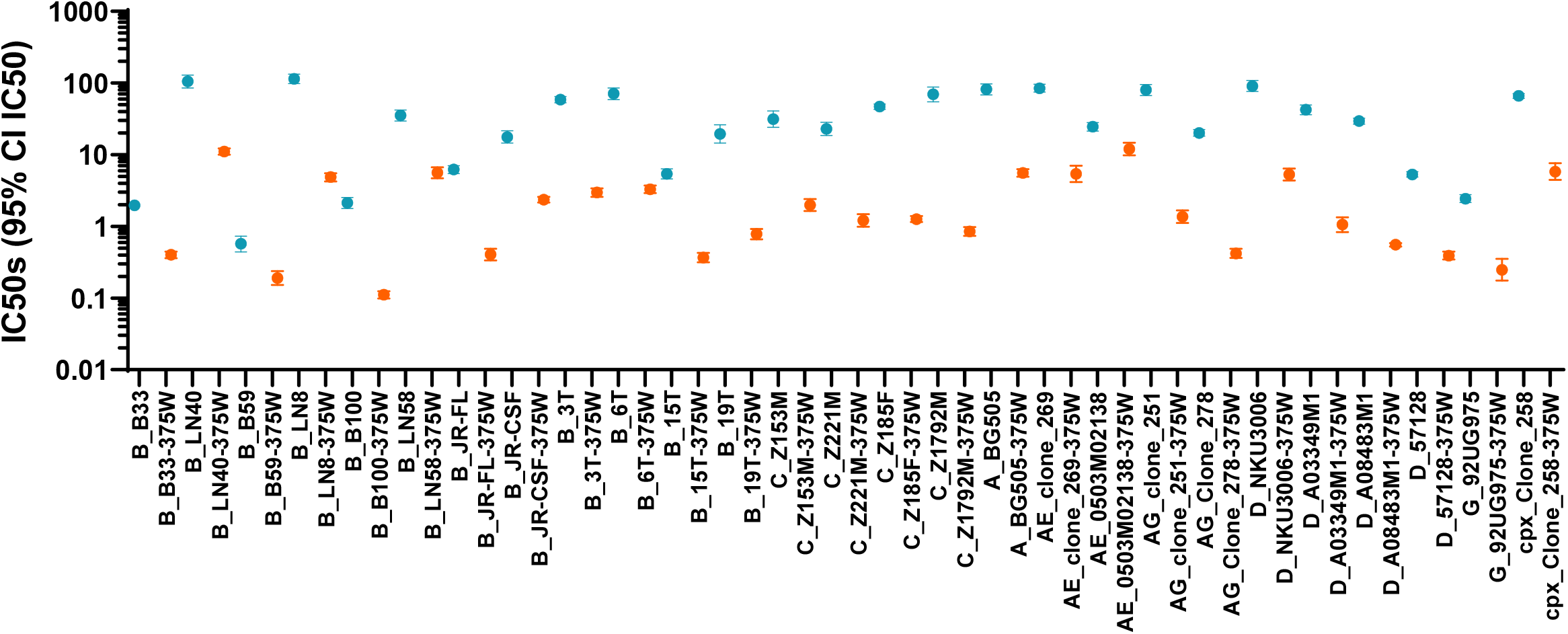
sCD4 neutralization. Geometric mean IC50s and 95% confident intervals were calculated and plotted in GraphPad. Two separate neutralization assays were performed, and each experiment was made in duplicate. The maximum concentration of sCD4 used to calculate IC50 was 400ug/ml. IC50s and confidence intervals for Wild type (WT) and 375W mutants are presented in blue 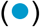or orange 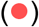circles and error bars, respectively.

**Figure 2:**
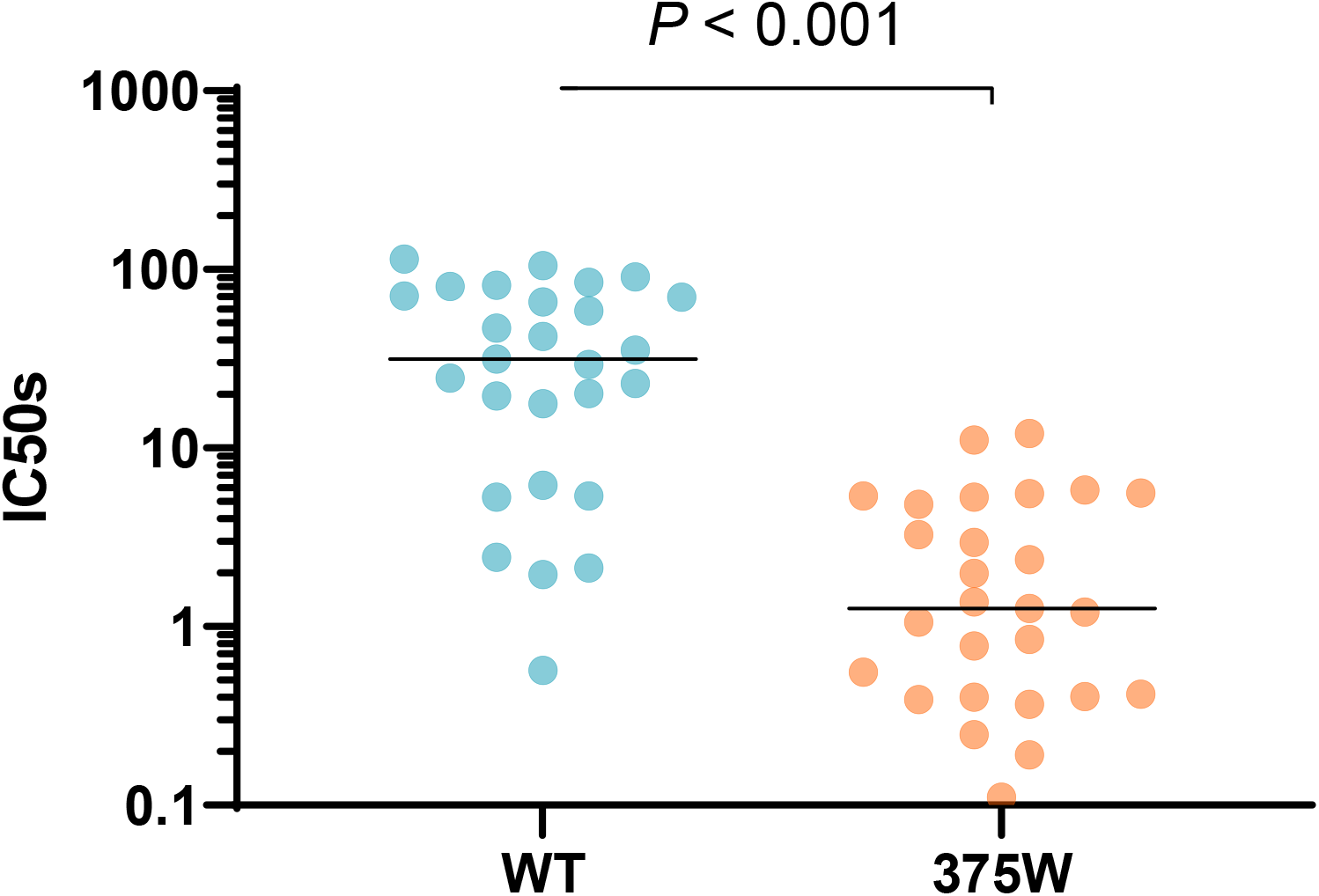
Compare the means of WT and 375W sCD4 neutralization IC50s from 2 experiments. 375W mutation decreases IC50 values compared with the WT Envs. The black line represents the median. WT and mutants were presented as light 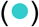and dark 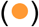gray circles. Analysis Wilcoxon test (paired nonparametric t-test) showed significant differences between WT and mutants IC50s (*p* value <0.001).

### 375W mutation changes tropism

We next evaluated whether the presence of a tryptophan in position 375influenced the biological property of tropism i.e. mac-tropism. Macrophages are cells that express low levels of CD4 receptor in the surface. When Envs with a better CD4 interaction encounter the receptor, the possibility that the strong binding effectively triggers fusion is higher. Then if 375W substitution increases CD4 binding, these mutants could infect better macrophages.We differentiated monocyte-derived macrophages (MDMs) from at least 3 healthy doors and infected with WT and 375W mutant pseudoviruses in duplicate wells. Each experiment was repeated at least 3 times and macrophage titers were normalized to those measured on TZM-bl titers (as described in the Methods section). Clade B and C (figure 3A) were tested together, while the other subtypes and CRF (figure 3B) were tested each other. Each assay included all the mac- and non-mac-tropic reference Envs used as positive and negative controls, respectively. The Central Nervous System (CNS)-derived Envs such as B33, B59, B100 and JR-FL are macrophage-tropic viruses that have been used extensively in our lab as positive controls for mac-tropism (1, 3). LN40, LN8, LN58 and JR-CSF are non-mac tropic viruses and were obtained from the same patients as B33, B59, B100 and JR-FL, respectively. For test Envs that were not derived from brain samples and which scored >1% macrophage titer (compared to TZM-bl) but that did not reach the macrophage titers of our brain mac-tropic reference Envs, we considered intermediate mac-tropic Envs.

**Figure 3:**
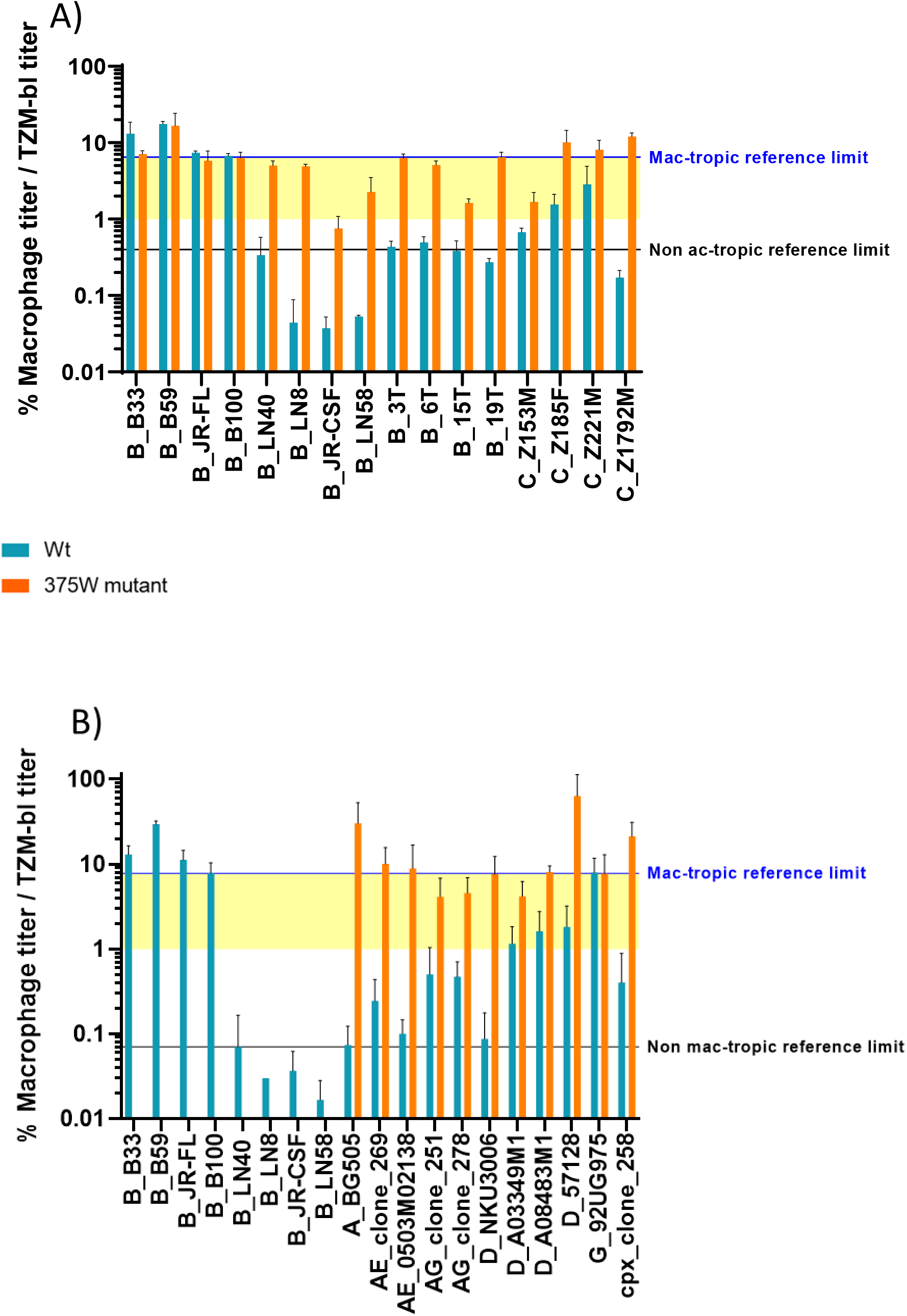
Percentage of macrophage titers vs TZM-bl titer. Each infection was repeated three times using differentiate MDM from at least 3 different healthy human blood donors. Titers in macrophages were normalized to TZM-bl titers. The data of subtype B and C (A), and other clades (B) are represented. Blue and orange bars represent WT and 375W mutant Envs, respectively. Black and blue line indicated higher titer of non-mac-tropic and the lower titer to consider that an Env is mac-tropic. In yellow, the region of intermediate mac-tropic Envs. SEM is represented on the top of each bar in black.

We found that the 375W mutation increased macrophage infectivity in all non mac-tropic and intermediate variants (figure 3A and B). C_Z185F, and C_Z221M wt Envs are defined as already having intermediate mac-tropic phenotypes. The 375W mutants of our non mac-tropic reference Envs (LN40, LN8, LN58 and JR-CSF) each infected macrophages much more efficiently than their WT counterparts. LN40, LN8 and LN58 each acquire intermediate mac-tropic phenotype when 375W substitution is present, as occurs with the other clade B Envs (3T, 6T, 15T and 19T) and clade C_Z153M. The presence of a tryptophan at position 375 in Z185F, Z221M and Z1792M Envs switched the phenotype to mac-tropic Envs while 3T and 19T mutants were borderline for mac-tropism (normalized macrophage titers of 3T and 19T are 6.33 and 6.49, respectively, compared with B100 that is 6.75).

We observed that the 375W mutants of the mac-tropic B33, B59 and JR-FL reference Envs showed reduced macrophage infectivity (Figure 3A) compared to their wt Envs. It is possible that these primary wt Envs have evolved to infect cells with low levels of CD4 receptor in brain and the introduction of the 375W mutation could alter an optimally adapted trimer structure, thus decreasing infectivity. Of note, the same mutation in the B100 mac-tropic Env also slightly decreases macrophage infectivity, consistent with this argument. Surprisingly, G_92UG975 Env is also mac-tropic Env with normalized macrophage titer of 7.92 close to B100 that is 7.81 (Figure 3B). Interestingly, G_92UG975 Env mutant showed the same slightly reduced macrophage infectivity pattern as for the B100 mac-tropic mutant.

We next tested subtype A, D and G, CRF and cpx wt and 375W variants separately and compared with the mac- and non mac-tropic references (Figure 3B). G (92UG975) and Clade D (A03349M1, A08483M1 and 57128) wt Envs showed mac-tropic and intermediate mac-tropic profile, respectively, indicating the presence of such mac-tropic and intermediate phenotypes outside of CNS (CSF and brain tissue) for these non B and C subtypes. The 375W mutation increased macrophage infection for these Envs as well as the CRF and cpx variants except for G_92UG975. Thus, A_BG505, AE_Clone_269, AE_0503M02138, D_A08483M1 and D_57128, and cpx_Clone_258 variants become mac-tropic when a tryptophan in position 375 is present. 375W D_NKU3006 and G_92UG975 mutants, exhibited intermediate phenotype but are borderline mac-tropic titers.

AE_0503M02138 was classified as X4 virus (22) and did does not infect macrophages well. However, it infects better than non-mac-tropic reference variants. When 375W substitution was introduced, an infectivity increase is observed changing to mac-tropic phenotype.

Finally, we compared WT and mutant Envs for macrophage titers adjusted to TZM-bl titers (see Materials and methods) and found that they are significantly different (p < 0.001) (Figure 4). In summary, we found mac-tropic and intermediate mac-tropic primary isolates outside the CNS from Clade G, and C and D, respectively. 373W induces an enhanced interaction with CD4 (Figure 1 and 2) and increases the macrophage infection of primary isolates from different subtypes and at different disease stages.

**Figure 4:**
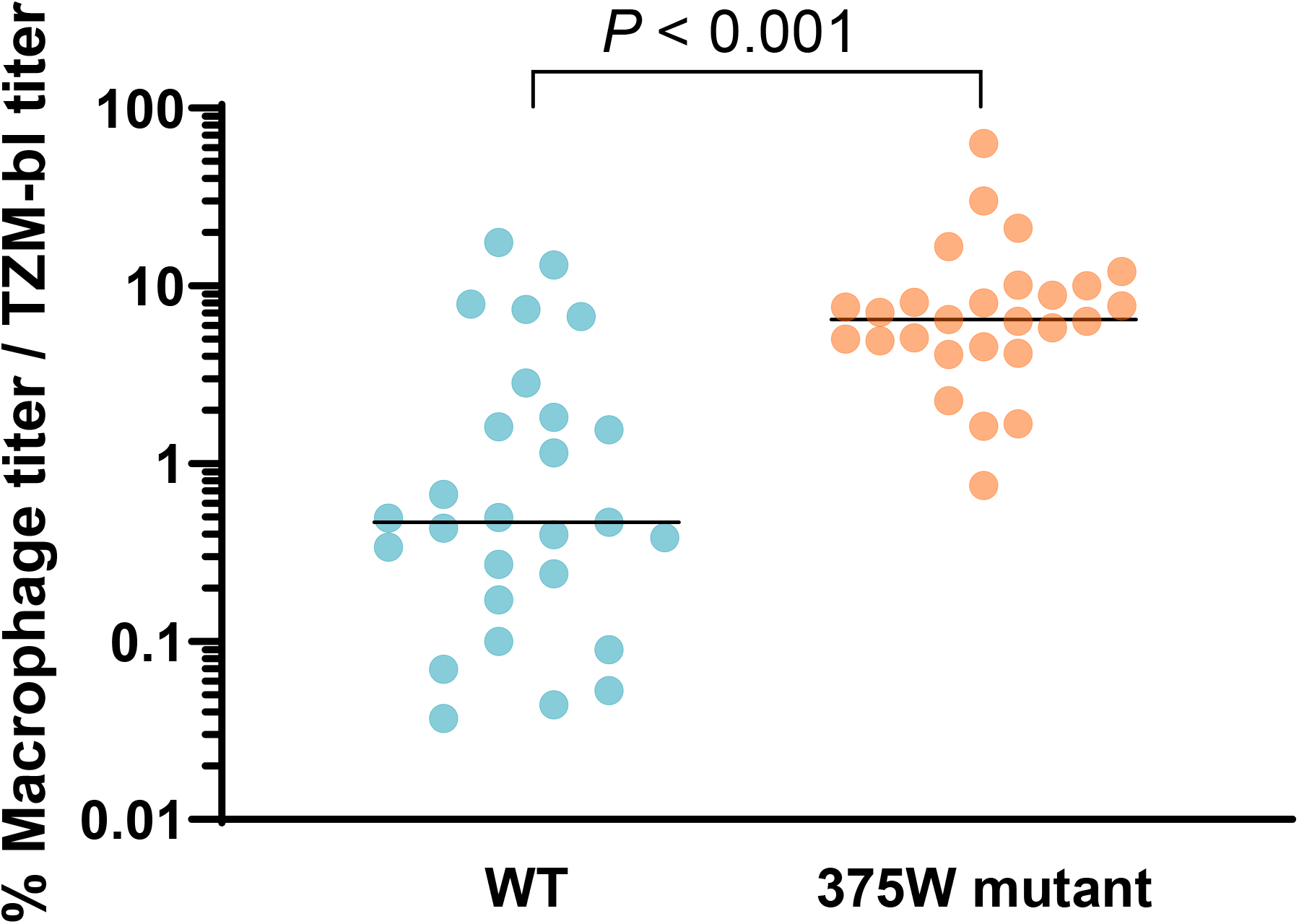
Comparison of the means of WT and 375W % macrophage titers/TZM-bl. The black lines represent the median. WT and mutant viruses were presented as blue 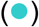and orange 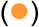circles, respectively. Analysis using Wilcoxon test (paired nonparametric t-test) showed significant differences between WT and mutants in % macrophage titers/TZM-bl (*p* < 0.001).

### Comparison of sCD4 neutralization IC50s and macrophage titers

We next determined if there is a correlation between mac-tropism and sCD4 neutralization. We compared macrophage titers with sCD4 IC50s (Figure 5). We used the Spearman correlation test to calculate the P value since the data were not normally distributed. We found that there was a strong correlation between mac-tropism and CD4 binding as shown by the P value < 0.001.

**Figure 5:**
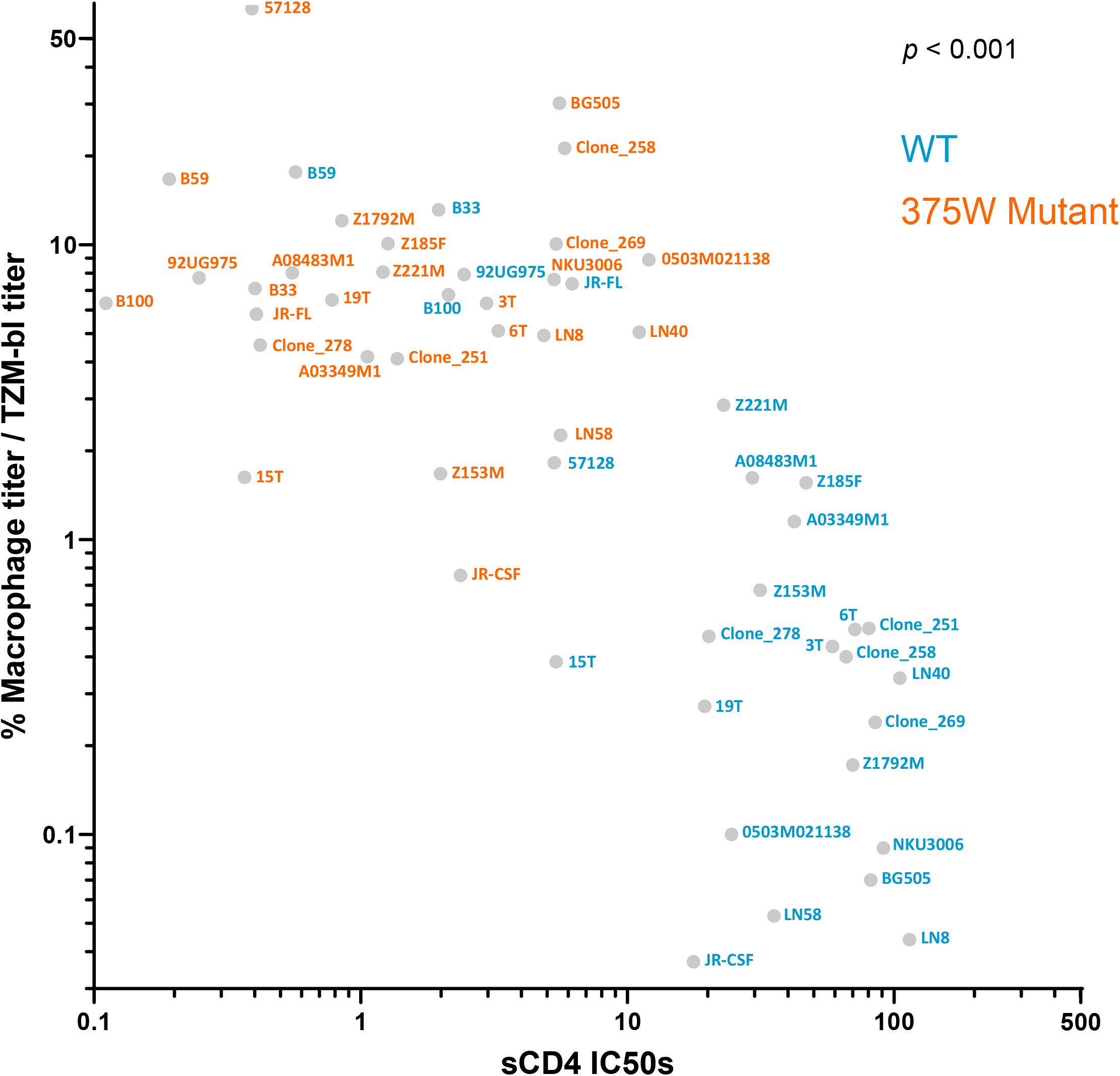
Correlation of % macrophage titers/TZM-bl vs sCD4 IC50 means. WT and 375W mutant were represented in blue and orange, respectively. The analysis of Spearman test (nonparametric correlation model) showed significant correlation between tropism and sCD4 neutralization (*p* < 0.001).

This relationship does not explain why B_B100 and G_92UG975 mutants have slightly less macrophage titers but the affinity for sCD4 of these mutants (measured by sCD4 inhibition) is 19 and 10 times higher compared with the wt, respectively. In addition, several mac-tropic (B_B33, B_B59, and B_JR-FL) Envs lost macrophage infectivity when 375W was introduced. Thus, B_B33, B_B59, B_JR-FL mutants show around 5, 3, and 15 increased sensitivity to sCD4 compared with the wt. These observations are clear indications that additional factors, apart from Env:CD4 interactions, influence mac-tropism.

### The effects of 375W on mAb epitopes

#### B6 and 17b neutralization

T-cell-line adapted (TCLA) or laboratory HIV-1 strains that have been passaged in the lab without the presence of human immunity evolve Envs that are more accessible to antibody binding, gp120 shedding and trimer opening to expose the CD4bs (23). On the contrary, primary isolates are subject to the immune pressures that select for Envs with closed trimers. The monoclonal antibody (mAb) b6 interacts with an epitope overlapping the CD4bs and binds clade B lab strains such as HXBc2 (24). b6 mAb binds to a very open CD4bs. We tested mAb b6 against subtype B and C primary isolates and it does not neutralize any these Env (data not shown) consistent with a CD4bs that is closed compared with the lab strains.

Following gp120 binding to CD4 there is a conformational change that exposes the coreceptor binding site and CD4-induced epitopes (CD4i). mAb 17b binds CD4i (25) including residues K121, R419, I420, K421, Q422 I423, and Y435 (26). We tested the clade B and C wt and 375W Envs described here and 17b in the absence of sCD4 does not neutralize any (data not shown). These results demonstrate that the 375W mutation does not induce a conformational change that exposes CD4i.

#### Neutralization by broadly active CD4bs mAbs

We were interested in understanding how the 375W mutation affects the exposure of CD4bs epitopes targeted by highly efficient bnAbs. To do so, we investigated b12, VRC01 and 3BNC117 mAbs in neutralization assays (Figure 6). VRC01 and 3BNC117 are mAbs that neutralize around 90% of the primary isolates from different clades.

**Figure 6:**
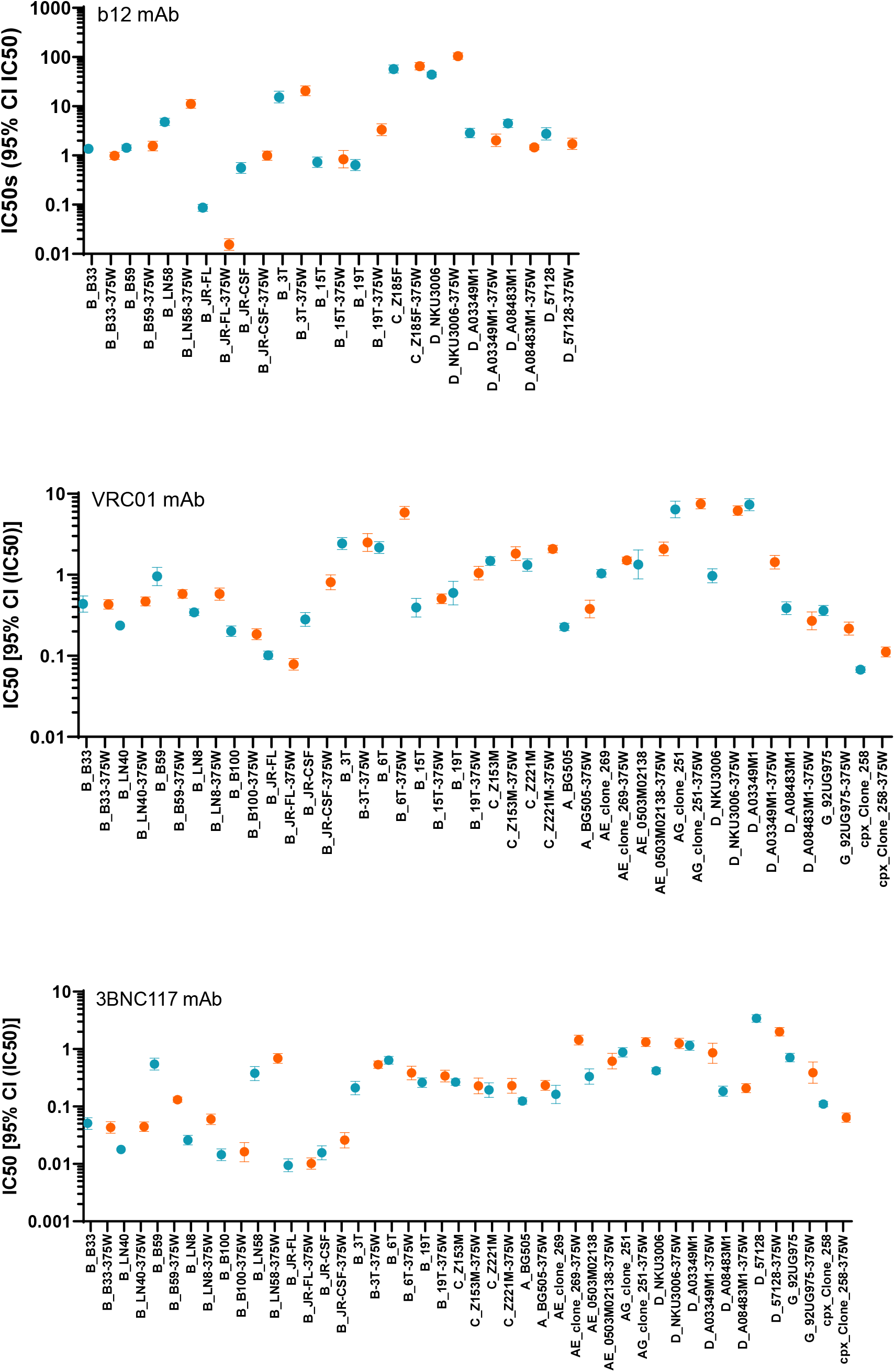
IC50 values of CD4bs mAb neutralization. Geometric mean IC50s and 95% confident intervals were calculated and plotted in GraphPad for b12 (A), VRC01 (B), and 3BNC117 (C) mAbs. Two different neutralization assays were performed, and each experiment was made in duplicate. Confidence intervals for Wild type (WT) and 375W mutants are presented in blue or orange error bars, respectively. WT IC50s are represented by blue squares 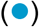and 375W mutant IC50s by orange circles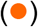.

##### b12 mAb

b12 mAb recognizes the CD4bs region but it is not a potent bNAb. Here, by using b12mAb, only clade B, C and D Envs were neutralized. Of note, clade A (BG505), B (B100, LN40, LN8, 6T), C (Z153M, Z221M, Z1792M), and G (92UG975), CRF_AE (clone_269 and 0503M02138), CRF_AE (clone 251 and 278) and cxp (clone 258) Envs were all resistant to b12 mAb (data not shown). Clade B LN40 is naturally resistant to b12 mAb due to the presence an asparagine at N386 together with an arginine at R373 which together block an Env pocket targeted by W100 on b12 (27).

There was no significant difference between wt and mutant 375W Env sensitivity to b12 neutralization (p = 0.22). Although, we did notice a tendency for higher IC50s when the 375W substitution was present in non-mac-tropic viruses indicating increased resistance (Fig. 6 A). Other researchers have also reported that the 375W mutation reduced b12 binding in gp120 monomers (ELISAs) and in neutralization assays (10, 11). Interestingly, in mac-tropic (B_B33, and B_JR-FL except for B59) and intermediate mac-tropic Envs (D_A03349M1, D_A08483M1, and D_57128 except for C_Z185F), the b12 interaction is slightly increased, while it is always decreased in non-mac-tropic Envs (Figure 6A). Non-mac-tropic primary isolates have a more closed CD4bs and the introduction of 375W substitution opens it. As consequence of this structural change, the b12 epitope may be altered, hampering b12 binding. The 375W conformational change resulting in more open mac-tropic and intermediate mac-tropic trimers improves or does not alter the b12 interaction. This could show that 375W residue induces slightly different conformational changes in the trimer depending on if the previous Env was more closed or more open. In summary, we have observed some differences between mac-tropic and intermediate vs non-mac-tropic CD4bs structure via b12 mAb neutralization.

##### The potent neutralizing VCR01 and 3BNC117 mAbs

VRC01 mAb potently neutralized pseudovirions carrying most wt Envs tested (Fig. 6B) but did not neutralize B_LN58, C_Z185F, C_Z1792M, AG_Clone_278 and D_57128. We observed that the presence of a tryptophan in position 375 slightly decreased VRC01 sensitivity for some Envs shown by small increases in the IC50 values (Fig. 6B). Despite these small variances, significant differences between wt and mutant VRC01 neutralization titers were observed (p=0.02). Interestingly, for mac-tropic (B_B59, B_JR-FL, B_B100, and G_92UG975), and intermediate mac-tropic Envs (D_A03349M1, and D_A08483M1), there is a neutralization shift of the 375W mutants towards sensitivity compared to WT Envs, except for mac-tropic B33 Env that did not change and for the intermediate mac-tropic Z221 Env neutralization sensitivity decreased. These results are consistent with a slight change of trimer structure in the CD4bs region for these mac-tropic and intermediate mac-tropic Envs since the 375W substitution did not affect non-mac-tropic mutants in the same way.

3BNC117 is a very potent and broad mAb since it neutralized nearly all Envs tested here at lower IC50 compared to the IC50s reached by VRC01 (Figure 6B and C). All Envs were neutralized with 3BNC117, except 15T, Z185F, Z1792M and Clone_278 Envs. Z185F, Z1792M, and Clone_278 Envs are also resistant to VRC01 bnAbs consistent with changes in the residues that affect both these bnAbs (Figure 6B and C). There were not significant differences between wt and mutant neutralization titers for 3BNC117 (p=0.41). However, we did find a tendency of slightly increased resistance to 3BNC117 mAb in 375W mutants (Figure 6C).

In summary, we have corroborated some differences between mac-tropic and intermediate vs non-mac-tropic CD4bs structure by CD4 mAb neutralization when position 375 has a tryptophan residue. However, the 375W mutation did not abrogate the binding of potent CD4 mAbs including VRC01 and 3BNC117 or b12 in any Env tested, leaving their epitopes accessible. Immunogens carrying 375W could therefore be used as a vaccine to induce CD4bs bNAbs.

#### Neutralization by V3 loop/glycan mAb

We previously reported that 375W confers a trimer conformation that opens the CD4bs without exposing the occluded GPGR V3 epitope that can potently induce non-neutralizing Abs such as 447-52D (9). In this study, we evaluated the effect of the 375W mutation on neutralization sensitivity to mAbs 10-1047 and PGT128 that recognize conserved V3 loop/V3 glycan epitopes in our panel of 27 diverse HIV-1s.

N332 glycan has been reported as a target for 10-1074, while N332 together with N156, N301 and N157 are targeted by the PGT121 mab family. PGT128 also interacts with N332, N156 and N301 (28, 29). 10-1074 has some minimal interactions with N156 and N301. N156 and N301 are present in all Envs investigated with the exception of AE_0503M02138 that lacks N301. However, B_B100, B_15T, C_Z153M, A_BG505, AE_0503M02138, D_NKU3006, and cpx_Clone_258 Envs do not have an asparagine at position 332 and were not neutralized by PGT128, nor by 10-1074 mAbs (Figure 7A). 10-1074 and PGT128 mAbs also bind the V3 peptide motif GDIR (28, 29). All Envs tested here have this motif except for B_6T, C_Z221M, C_Z1792M and D_NKU3006 that have GNIR. Nevertheless, these latter Envs are neutralized by both mAbs with the exception of D_NKU3006 that also lacks the N332 glycan (fig. 7A and B).

**Figure 7:**
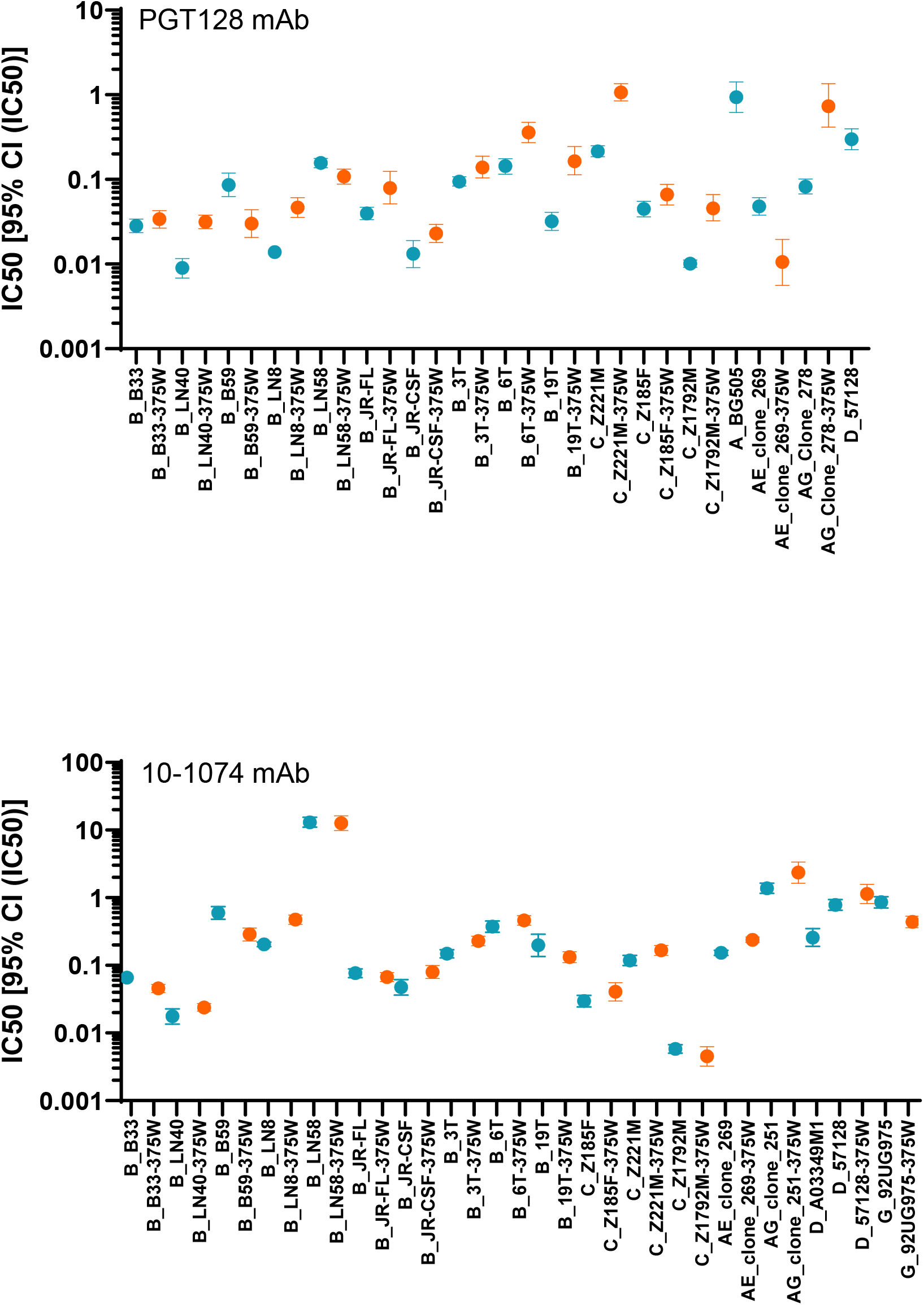
IC50 values of V3 loop/glycan mAb neutralization. Geometric mean IC50s and 95% confident intervals were calculated and plotted in GraphPad for PGT128 (A), and 10-1074 (B) mAbs. Two different neutralization assays were performed, and each experiment was made in duplicate. Confidence intervals for Wild type (WT) and 375W mutants are presented in blue or orange error bars, respectively. WT IC50s are represented by blue squares 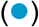and 375W mutant IC50s by orange circles 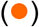.

When a tryptophan at position 375 is present, we found that mac-tropic (B_B33, B_B59, B_JR-FL, and G_92UG975) increased binding for 10-1074 mAb (Figure 7B). It could indicate 375W substitution in the mac-tropic variants with more open CD4bs modify the apex of the trimer facilitating the interaction of 10-1074 mAb. However, this IC50 reduction was also observed in three non-mac-tropic Envs including B_LN58, B_19T and C_Z1792M indicating that there is not a clear neutralization pattern associated to mac-tropism. For 375W D_A03349M1 mutant, IC50 could not be calculated at 100ug/ml of 10-1074 mAb (data not shown). The same is true for A_BG505 and D_57128 mutants when Envs were neutralized with PGT128 mAb at 200ug/ml (data not shown). The same maximum concentrations were used to neutralize and calculate A_BG505, D_A03349M1, and D_57128 wt IC50s. The 375W substitution could potentially induce a glycan movement causing a partial or completely loss of mAb interaction with the Env. For some Envs, it is possible that an increase of mAbs concentration allow the calculation of the IC50. For the PGT128 mAb, 375W mutants are less or more sensitive compared to wt depending on the Env tested. IC50 values do not show a significant difference between wt and mutants for 10-1074 and PGT128 mAbs with p=0.49 and p=0.07, respectively.

Overall, these mAbs interact better with clade B and C compared with other subtypes, CRF and cpx in our panel. Our results indicate the 375W mutants retain sensitivity to broadly neutralizing V3 loop/glycan 10-1047 and PGT128 mAbs. Thus, 375W Env mutants that carry an exposed CD4bs (as we previously reported), consistently maintain exposure of the vaccine-desirable, glycan dependent, V3 loop epitopes targeted by highly potent bnAbs.

#### Neutralization by V1V2 loop/glycan mAb

We next tested our primary Env panel against PGT145, PG9 and PG16 (Figure 8), less potent bnAbs that recognize N-linked glycosylation sites such as N160 and N156 and sometimes N173 as a compensatory residue of V2 loops in the trimer apex (30). PGT145 does not directly interact with N156 but the loss of that glycan abrogates its interaction with the trimer apex. N156 is present in all Envs tested. However, none of our Envs has an asparagine at position 173. N160 glycan is not present in B_LN40, B_15T, C_Z185F, D_NKU3006, and G_ 92UG975 Envs. None of these primary isolates were neutralized by PGT145, PG9 or PG16 mAb (data not shown).

**Figure 8:**
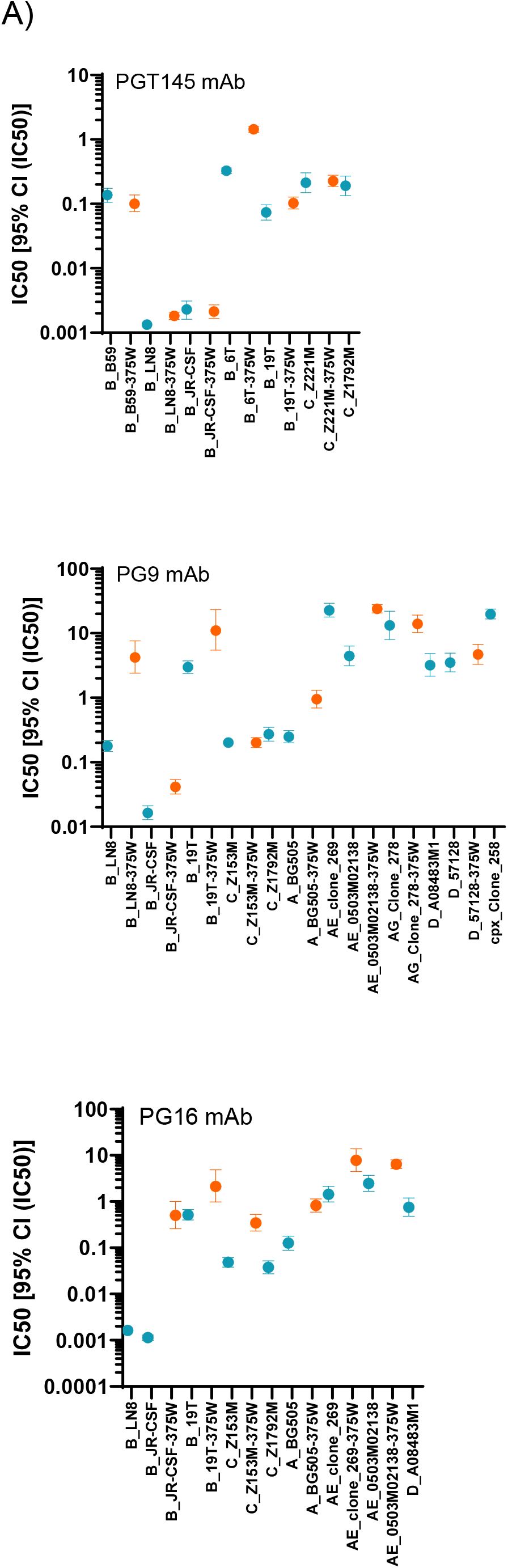
IC50 values of V1V2 loop/glycan mAb neutralization. Geometric mean IC50s and 95% confident intervals were calculated and plotted in GraphPad for PGT145 (A), PG9 (B), and PG16 (C) mAbs. Two different neutralization assays were performed, and each experiment was made in duplicate. Confidence intervals for Wild type (WT) and 375W mutants are presented in blue or orange error bars, respectively. WT IC50s are represented by blue squares 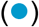and 375W mutant IC50s by orange circles 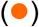.

PGT145 targets a quaternary epitope simultaneously interacting with the 3 gp120 monomers in pre-fusion conformation (30). 3BNC117 mAb binding to the CD4bs was reported to increase the space between N160 residues of each of the 3 monomers and enhance the accessibility of PGT145 epitope (30). IC50s were not calculated in non-subtype B and C panel, since these Envs do not bind PGT145. K121A and R166A substitutions have also been associated with a reduction in PGT145 neutralization (30). In our panel, all Envs tested have R166 except for B_3T, B_15T, C_Z185F, AG_clone_251 and _278. For K121, C_Z153M has T and AE_0503M02138 has I at this position. All these primary Envs are resistant to PGT145 mAb. Data obtained from Envs that are sensitive to PGT145 mAb indicates that there are no changes or marginally increased resistance to this mAb for 375W mutants (Figure 8A). However, the differences between wts and 375W mutants are not statistically significant (p=0.44).

PG9 and PG16 bind gp120 monomers as well as the trimer, while PGT145 only binds quaternary Env structures (31). However, PG16 preferentially binds to trimers while PG9 binds two of the three gp120 protomers in the trimer (32). These differences could explain why PG9 and PG16 do not always neutralize the same Envs in our panel. C_Z153M mutant has the same PG9 IC50 value as the wt indicating that the 375W substitution does not affect the interaction with this mAb. In general, a tryptophan at position 375 increased resistance to these less potent bnAbs (Figure 8B and C). Differences between WT and 375W mutants are significant with p= 0.0078 and p= 0.03 for PG9 and PG16 mAb, respectively. However, few of the Envs tested were sensitive to these mAbs, although it is possible that the 375W substitution alters the apex structure of the trimer.

IC50s could not be calculated for C_Z1792M mutant using PGT145 where we used up to 400ug/ml of mab, or for B_LN8, C_Z1792M and D_A08483M1 mutants using PG16 concentrations also of up to 400ug/ml, and C_Z1792M, AE_Clone_269, D_A08483M1, and cpx_Clone_258 mutant using PG9 concentrations up to 400ug/ml (data not shown). The C_Z1792M mutant was resistant to all V1V2 loop/glycan mAbs, and D_A08483M1 mutant was resistant to PG9 and PG16. The maximum mAb concentrations needed to reach 100% of neutralization for C_Z1792M wt Env were lower (100ug/ml for PGT145, 2ug/ml for PG16, and 5ug /ml for PG9) showing that C_ Z1792M mutant is resistant to these mAbs. 400ug/ml of PGT145, PG9 and PG16 could not neutralize this 375W mutant when the wt was neutralized at lower mAb concentration values. To calculate the IC50 for D_A08483M1 wt we used 400ug/ml of PG9 and PG16 mAbs as maximum concentrations. At these levels of ab, D_A08483M1 wt was weakly sensitive to these mAbs and the 375W mutant was resistant. These V1V2 loop/glycan mAbs neutralize fewer Envs of our panel (Figure 8) and are less potent compared to the CD4bs and V3 loop/glycan mAbs. Moreover, there is a loss of affinity for PG9 and PG16 mAbs when 375W is introduced.

## Discussion

We previously reported that the 375W mutation opens the HIV trimer and facilitates binding to the CD4 receptor without exposing the V3 loop (9). HIV trimers carrying 375W could therefore be good vaccine candidates to develop Abs against the CD4bs. The 375W mediated increase in sCD4 binding was previously observed for clade B,YU2 too (10) but also for clade A, B, C and D SHIVs (16). These findings support our results that 375W residue increases CD4 interactions in all Env tested. In this study, we also show that the 375W substitution has biological significance by increasing macrophage infection of many our Envs in our panel. However, and of note, the 375W mutation actually reduced macrophage infectivity for mac-tropic primary isolates, particularly those highly adapted for replication in brain macrophages. These latter results are also consistent with previous data showing that 375W reduced infectivity for mac-tropic clade B, YU2 (10, 11, 33).

The ideal HIV immunogens will likely present the CD4bs in an appropriate conformation to induce bnAbs that approach the CD4bs at an optimal angle (34). Such immunogens will be presented as Env trimers and will be designed to limit induction of ‘off-target” non-nAbs especially against CD4i and V3 loop epitopes. Trimer based immunogens have been developed in the form of BG505 SOSIP.664. However, this immunogen induced the production of non-nAbs against the trimer base (7, 35). Furthermore, CD4 binding induces conformational changes in BG505 SOSIP.664 that lead to the exposure of non-neutralizing epitopes (CD4i and V3 loop) and this seems to occur during immunization (35). Env neutralization phenotypes are classified as tier 1A and B, 2 and 3 depending on the numbers of epitopes exposed and neutralization sensitivity. In tier 1, Envs that have open trimers were classified as tier 1A while Envs with a more intermediate conformation are defined as tier 1B. Together, these Envs expose a higher number of epitopes and are easier to neutralize. In contrast, tier 2 and 3 Envs with more closed trimer conformations expose less epitopes and are more resistant to neutralization (36). Further studies showed that the derivative, DS-SOSIP.4mut, modified to stabilize the SOSIP trimer in a more prefusion and closed conformation, reduced CD4 binding and a V3-sensitive tier-1 response while increasing tier-2 antigenicity (6). However, this construct did not induce bnAbs

In single molecule fluorescence resonance energy transfer (smFRET) experiments, different conformational states of Env were observed e.g. state 1 (a closed conformation), state 2 (asymmetric intermediate conformations with one protomer open) and state 3 (asymmetric open conformation with three protomers open that are bound to three CD4 molecules). The binding of a single CD4 molecule into one protomers of the trimer induced a state 3 conformation, while the other protomers (without CD4 bound) transitioned to state 2 (37). The BG505 SOSIP.664 trimers predominantly stay in state 2 in smFRET (38) while primary isolate trimers exhibit state 1 conformations (38, 39), indicating that they have different native structures. SOSIP immunogens have been modified to prefusion-closed structures using introduced disulfide bonds. This approach may help modulate the trimer structure, optimizing the presentation of desired epitopes and the capacity to enhance Ab breath in responses against the trimer. The introduction of 2 disulfide bonds in BG505 (non-mac-tropic Env) and JR-FL (mac-tropic Env) Envs stabilized the so-called state 2 conformation as identified using smFRET (38), while such Envs are usually predominantly in a state 1 conformation. These novel approaches may eventually result in the identification of trimers that are immunogenic for bnAbs. We envisage that 375W will play an important role in the development of such immunogens.

We propose that HIV-1 375W Envs will be good candidates for vaccines aiming to elicit bnAbs targeting the CD4bs. Since VRC01 and 3BNC117 neutralize 375W mutants as efficiently as wt, it indicates that 375W immunogens carry these highly desirable epitopes for vaccines and would have the potential to generate bnAbs maintain the appropriate angle of approach to the CD4bs. Our data also indicate that 375W Envs retain V3/glycan epitopes (e.g targeted by PGT128 and 10-1074 mabs) exposed on mutant primary viruses of different clades, CRF and cpx. Such Envs may therefore have the potential to be immunogens for bnAbs targeting such glycans. In addition, in a similar strategy where specific glycans are removed from Env immunogens to enhance induction of Abs against the CD4bs (40), 375W immunogens could be used as a first boost to present the CD4bs, before being followed by consecutive boosts with diverse, prefusion-closed Env trimers. This approach would represent an attempt to mimick the induction of bnAbs in chronic infection. Our data also highlight another advantage of the 375W mutation for vaccines, in that it exposes the CD4bs and introduces the similar conformational change in all clades, CRFs, and cpx as well as in Envs from different stages of infection tested. Recently, a novel HIV-1 mRNA vaccine was tested in rhesus macaques and heterologous tier-2 SHIV responses were observed (41). This new vaccine platform has the advantage that can present different immunogens in their natural conformations at the same time and would be highly appropriate for the inclusion of W375 trimer immunogens.

We found that bnAbs such as VRC01, and 3BNC117 neutralized almost all wt Envs we tested, while 10-1074 and PGT128 are more effective neutralizing non-clade B and C variants. For VRC01 some significant IC50 differences were observed (p=0.02) between wt and mutant Envs. Thus, there is a tendency that the presence of a tryptophan in position 375 slightly decreased VRC01 sensitivity in non-mac-tropic variants, however, this did not ever abrogate VRC01 interaction. Interestingly, almost all mac-tropic, and intermediate mac-tropic 375W Envs were neutralized with increased sensitivity for VRC01 mAb. 375W substitution also altered the b12 mAb interaction. In mac-tropic and intermediate mac-tropic variants, the 375W residue induced a slightly better exposure of b12 epitope compared with non-mac-tropic variants. It is possible that the structural change induced by 375W substitution alters the approach angle of these mAbs (42) causing different effects depending on how each Env is (or isn’t) adapted for mac-tropism. No significant differences in the accessibility to 3BNC117, 10-1074 and PGT128 epitopes between variants with different tropisms were observed when the 735W substitution was present. These results show that small structural changes in the trimer can be detected between mac-tropic, intermediate and non-mac-tropic Envs using certain mAbs in neutralization assays.

We and others have reported on highly mac-tropic Envs derived from brain tissue, particularly in subjects with neuro-AIDS (3). Based on our results here, we were able to define Envs from clade C, and D, with an intermediate mac-tropic phenotype, that have the capacity to infect macrophages at lower levels compared with highly mac-tropic Env. In children infected with clade C, variants were detected that could infect Affinofile cells via low CD4 at modest levels suggesting that they could be precursors of mac-tropic viruses (2). Subtype differences in the capacity to induce HIV-associated cognitive impairments in Uganda and Ethiopia suggest different biological characteristics between clades (43). For example, in Clade D infected patients, it is more common to develop HIV-1 dementia compared with clade A infected individuals (44, 45). However, individuals infected with clade D have been reported to progress to AIDS more frequently compared with clade A (43, 46). Here, we found that 3 of 4 clade D Env have an intermediate mac-tropic phenotype. HIV-related neurocognitive impairments have also been reported in patients infected with clade G in Nigeria (47). Interestingly, our clade G Env (not from CNS tissue) was mac-tropic. In South Africa, it has been found that some subtype C viruses induce cognitive impairments independent of a tat polymorphism previously associated with neurocognitive impairments (48). The clade C Envs (supplied by Dr. Derdeyn and Dr. Hunter) used here come from Zambia in South Africa and these also show an intermediate phenotype (49, 50). It’s possible that such intermediate mac-tropic variants circulating in blood could invade the brain and quickly adapt to the environment there, increasing mac-tropism and the likelihood of HIV-1 dementia. An alternative explanation is that these intermediate mac-tropic viruses could be an adaptation of mac-tropic viruses released from brain to blood where immune pressure is higher.

It is likely that the stage of disease influences the presence of different Env phenotypes. All T/F primary isolates including A_BG505, C_Z1792M and clade B (3T, 6T, 15T, and 19T) are non-mac-tropic. It has been described that mac-tropic viruses are not favorably transmitted in ectocervical tissue (51). Interestingly, some of Envs obtained from the acute phase showed intermediate mac-topic phenotypes (Clade C Z185F, and Z221M). In the acute stage, HIV-1 is at a high peak of replication and many variants are generated in the viral population. It is easy to envisage that these will include intermediate mac-tropic variants that could invade the brain establishing the infection in the CNS.

Surprisingly, a tryptophan at position 375 reduced macrophage infection in clade B Envs derived from brain tissue, although B_B100 was slightly less affected than G_92UG975 mac-tropic Env from cultured PBMC DNA. We could deduce that these mac-tropic Envs carry trimers are highly adapted to the brain environment and carry a succession of Env substitutions that confer this phenotype. Thus, the 375W substitution may have a negative impact on such optimally mac-tropic Envs. Nevertheless, previous studies indicate that a single mutation can transform a non-mac-tropic virus to mac-tropic (52-54). Here, we therefore observed the same effect in subtype A, C, CRF01_AE and CRF13_cpx non-mac-tropic as well as in intermediate clade D variants when 375W substitution is introduced. Moreover, an X4 CRF01_AE (0503M02138) also conferred a mac-tropic phenotype when 375W was introduced. If enhanced exposure of the CD4bs evolves in the blood or peripheral tissues, such Envs could be easier to neutralize by nAbs and this pressure likely reduces the presence of such variants there. Nevertheless, here we describe mac-tropic and intermediate mac-tropic viruses outside the CNS from different clades. The envelope structure and exposure of the CD4bs of such variants is unknown but could explain why Clade D infected individuals are more likely to develop dementia. More research is needed to understand the role of mac-tropism in peripheral tissues and to study mac-tropism in non-clade B Envs.

In summary, we found that the 375W substitution increased CD4 binding in all Envs tested from different disease stages. This increase in affinity correlated with an improvement of the macrophage infection with the exception of mac-tropic Envs which were already highly adapted to exploited low levels of CD4 for entry. Intermediate mac-tropic Envs were defined in this paper and found to be present outside the CNS perhaps potentially showing that these Env are present before brain infection. The presence of mac-tropic Env in blood indicate that mac-tropic Envs initially evolve outside the CNS or are sometimes released from there. Finally, 375W mutation could be a good substitution to introduce into immunogens for the development of vaccines against CD4bs.

## Materials and methods

### Cells and reagents

293T/17 cells (ATCC CRL-11268) (57) were used for transfection of DNA plasmid constructs to prepare HIV-1 Env+ pseudovirions. HeLa TZM-bl cells from the HIV Reagent Program were used in to titrate pseudovirions and for neutralization assays (1). HeLa TZM-bl carry β-galactosidase and luciferase reporter genes under the control of an HIV long terminal repeat. They express high levels of CD4, and CCR5, and are susceptible to infection by both R5 and X4 HIV-1 isolates (56, 70).

Soluble CD4 (sCD4) was from Progenics. MAbs b12, 2G12, PG9, and PGT145 were from HIV Reagent Program and Polymun Scientific. 17b, 447-52D, VRC01, PG16, PGT128, 3BNC117 and 10-1074 mAbs were from the HIV Reagent Program. b6 was provided by Dr. Dennis Burton, Scripps, La Jolla, CA.

### Env primary isolates

Expression plasmids carrying Envelopes (Envs) derived from molecular clones of transmitted/founder (T/F), acute stage, late disease stage macrophage-tropic and non-macrophage-tropic envelopes were provided by the HIV Reagent Program, Dr. Jonathan Ball (3), University of Nottingham, Nottingham, UK, Dr. Cynthia Derdeyn and Dr. Eric Hunter, Emory University, Atlanta, GA (62) or isolated in our laboratory (1). These Env+ plasmids were used for pseudovirus production (Table 1).

Envelope genes from transmitted/founder (T/F) (3T, 6T, 15T, and 19T, and Z1792M), acute stage (Z153M, Z221M, and Z185F), chronic infection M-tropic (B33, B59, JR-FL, B100), non-M-tropic (LN40, LN8, JR-CSF and LN58) primary isolates were previously inserted in pSVIIIenv.

### Direct mutagenesis

375W substitutions in all the primary isolates were introduced by site-directed mutagenesis in plasmids carrying Envs using a QuikChange site-directed mutagenesis kit (Stratagene Inc.) and mutagenic primers to introduce each mutation (56). 375W mutants of all the primary isolate Envs were obtained by direct mutagenesis in their original plasmids. The gp160 insert was sequenced to confirm the presence of the 375W mutation and the absence of any undesired mutation formed during the direct mutagenesis PCR.

### Env pseudovirions production and titration

Calcium phosphate was used to cotransfect plasmids carrying wild type (WT) or 375W mutant envs with pNL4.3 ΔEnv that does not express Env into 293T cells. This latter plasmid has a 2 amino acid insertion changing the reading frame in Env. Env+ pseudovirions from the cell supernatants were harvested 48 h after transfection, clarified (1,000 x*G* for 10 min), aliquoted, and stored at -180°C.

WT and mutant pseudoviruses were titrated in HeLa TZM-bl as described previously (1, 56, 57). Infections were performed in duplicate. Briefly, serially diluted (10-fold dilutions) cell-free virus supernatant was incubated with HeLa TZM-bl. Infected cells were fixed 2 days post-infection in phosphate-buffered saline (PBS)-0.5% glutaraldehyde at 4°C and stained with X-Gal (0.5 mg of 5-bromo-4-chloro-3-indolyl-β-D-galactopyranoside/ml [Fisher Bioreagents Inc.] in PBS containing 3 mM potassium ferrocyanide, 3 mM potassium ferricyanide, 1 mM magnesium chloride for 3 h. Focus-forming units (FFU) were estimated by counting individual or small groups of blue-stained, infected cells and the numbers of FFUs/ml were then calculated.

### Inhibition and Neutralization assays

Neutralization assays with carried out using the panel of mAbs in table 2 and inhibition assays with soluble CD4 (sCD4) were performed by reduction in FFU following infection of HeLa TZM-BL cells with 200 FFU of wt or mutant Env+ pseudoviruses as described previously (57). Two individual neutralization or inhibition experiments were performed in duplicate wells. Positive controls included untreated Env+ pseudovirions and the negative controls, non-infected cells. IC50s were then calculated (see IC50 determinations and statistics).

**Table 2:** mAbs used in this study.

| Reagent | Epitope | References |
| --- | --- | --- |
| b6 | CD4bs | (24) |
| b12 |  | (65) |
| VRC01 |  | (66) |
| 3BNC117 |  | (67) |
| PG9 | Trimer association domain (TAD) V2q | (68) |
| PG16 |  | (68) |
| PGT145 |  | (69) |
| PGT128 | V3 loop/glycan | (69) |
| 10-1074 |  | (67) |
| 17b | CD4 induced (CD4i) | (25) |

### Infection of healthy donor macrophages

Human monocytes-derived macrophages (MDM) were obtained from healthy human blood donors after preparing blood monocyte monolayers and culturing for 2 days in medium containing macrophage colony-stimulating factor (R&D Systems) as described previously (56, 71). Versene was used to wash macrophages, three times while incubating at 37°C for 10 min to loosen cell attachments the day prior to infection. Macrophages were then gently scraped off and reseeded into 48-well tissue culture trays.

1.25 ×10^5^ MDMs in 48-well tissue culture trays were treated with DEAE dextran (56). Serial 10-fold virus supernatant dilutions were added, spinoculated and incubated for 2 h 15min at 37°C. Virus supernatant was rinsed twice with DMEM. After one week post infection, macrophages were washed once in PBS, then fixed in cold Methanol/Acetone and stained with X-Gal. The numbers of focus-forming units (FFUs) /ml were estimated (56). Three individual infections were performed in duplicate wells. Error bars in figures were calculated from replicate wells for individual experiments.

Previously, we reported that there is a range of macrophage infectivity in Envs derived from brain samples [3]. Considering these data together with the reference viruses introduced in each assay as controls, we establish that the percentage of macrophage titer (titer normalized respect to TZM-bl titer) < 1% represent non mac-tropic viruses while >1% viruses show a good infection in macrophages. The lower value of the mac-tropic references after repeat the experiment 3 times represent the limit considered to define a mac-tropic Env. Envs that show % macrophage titer >1% and lower than the lowest brain mac-tropic reference macrophage titer were considered intermediate mac-tropic Envs.

### IC50 determinations and statistics

GraphPad prism software (vX9.3.3) was used to analyze the data. Normalized data was evaluated in Excel and transformed in GraphPad. Sy.x and AICc were used to establish the goodness of fit and to decide the more appropriate model to calculate geometric mean IC50 and 95% CI of IC50. The nonlinear regression method used was a dose-response inhibition model.

Before applying a parametric or non-parametric test, we assessed if our samples are normally distributed using statistical tests or graphical methods. We applied two normality tests: 1)The Shapiro–Wilk test to calculate if normality was used because it is recommended and a more appropriate method for small sample sizes of less than 50 (72, 73) but you need to have unique values, and 2) Anderson-Darling test (the method of Royston) that keeps in mind the skewness and kurtosis to calculate how far the distribution is from asymmetry and shape Gaussian. Q-Q, frequency (histograms), violin, and box plots were used to corroborate the results of the two normality tests.

If our data was Gaussian distribution a paired t test was used to determine significant differences between the means of two groups of data, if not we applied two-tailed non-parametric Wilcoxon signed-rank test. We used an F test to decide if both normal distributed populations have the same standard deviation (SD), and then applied unpaired t test or t test Welch’s correction. If the data is not normally distributed, two-tailed, nonparametric Mann-Whitney test (Wilcoxon rank sum test) is used. To test correlations, we used Spearman nonparametric correlation or Pearson correlation if the data follow a Gaussian distribution. P value asterisks as were as New England Journal of Medicine (NEJM) guidelines [<0.033 (*), <0.002 (**), <0.001 (***)].

## Acknowledgments

This study was supported by NIH R01 Grants MH115773 and NIH R21 AI127231.

We thank Dr. Cynthia Derdeyn and Dr. Eric Hunter for providing HIV-1 clade C envelope clones Z1792MPL18DEC07.3G7Env, Z185FPB24AUG02ENV3.1, Z153MPL13MAR02ENV5.1, and Z221MPL7MAR03ENV2.1, Dr. George Lewis for mAb, 17b, Dr. Richard Burton for b6 and Dr. Jonathan K Ball for E21B100 and E21LN58 Envs.

We acknowledge and thank NIH HIV Reagent Program, Division of AIDS, NIAID, NIH for p1054.TC4.1499, p63358.p3.4013, p700010040.C9.4520 and pPRB958_06.TB1.4305 contributed by Drs. Beatrice H. Hahn, Brandon F. Keele, George M. Shaw; BG505-W6M-ENVC2 contributed by Dr. Julie Overbaugh; clones 251, 258, 269 and 278 contributed by Drs. D. Ellenberger B. Li, M. Callahan, and S. Butera; pCRII_92UG975.10 contributed by Dr. Feng Gao and Dr. Beatrice Hahn; 0503M02138.EC1, NKU3006.EC1, A03349M1, A08483M1.VRC09A and 57128.VRC15 contributed by Dr. Sodsai Tovanabutra; 3BNC117 and 10-1074 m Abs contributed by Dr. Michel Nussenzweig; b12 contributed by Dr. Dr. Dennis Burton and Dr. Carlos Barbas; VRC01 contributed by Dr. John Mascola; 447-52D contributed by Dr. Zolla-Pazner; 2G12 contributed by DAIDS/NIAID; PG9, PG16, PGT145 and PGT128 contributed by International AIDS Vaccine Initiative; contributed by ; 17b contributed by Dr. James E. Robinson; TZM-bl contributed by Dr. John C. Kappes and Dr. Xiaoyun Wu.

